# Micelle-like clusters in phase-separated Nanog condensates: A molecular simulation study

**DOI:** 10.1101/2023.05.18.541260

**Authors:** Azuki Mizutani, Cheng Tan, Yuji Sugita, Shoji Takada

## Abstract

The phase separation model for transcription suggests that transcription factors (TFs), coactivators, and RNA polymerases form biomolecular condensates around active gene loci and regulate transcription. However, the structural details of condensates remain elusive. In this study, for Nanog, a master TF in mammalian embryonic stem cells known to form protein condensates *in vitro*, we examined protein structures in the condensates using residue-level coarse-grained molecular simulations. Human Nanog formed micelle-like clusters in the condensate. In the micelle-like cluster, the C-terminal disordered domains, including the tryptophan repeat (WR) regions, interacted with each other near the cluster center primarily via hydrophobic interaction. In contrast, hydrophilic disordered N-terminal and DNA-binding domains were exposed on the surface of the clusters. Electrostatic attractions of these surface residues were responsible for bridging multiple micelle-like structures in the condensate. The micelle-like structure and condensate were dynamic and liquid-like. Mutation of tryptophan residues in the WR region which was implicated to be important for a Nanog function resulted in dissolution of the Nanog condensate. Finally, to examine the impact of Nanog cluster to DNA, we added DNA fragments to the Nanog condensate. Nanog DNA-binding domains exposed to the surface of the micelle-like cluster could recruit more than one DNA fragments, making DNA-DNA distance shorter.

**Author summary:** In eukaryotic transcription regulation, enhancer elements far from the promoter are known to modulate transcriptional activity, but the molecular mechanism remains elusive. One of these models, the phase separation model, suggests that transcription factors (TFs), coactivators, and transcription machinery form biomolecular condensates that include enhancer and promoter elements, making enhancer-promoter communication possible via the protein network in the condensate. Nanog, one of the core TFs involved in mammalian embryonic stem cells, has been reported to have the ability to form condensates. In this study, we addressed the structural details of Nanog condensates by performing residue-level coarse-grained molecular simulations. We found that Nanog formed micelle-like clusters via primarily hydrophobic interactions between tryptophan repeat regions in the C-terminal disordered domains within the condensate. On the other hand, highly charged N-terminal and DNA-binding domains were exposed to the surface of the micelle and were responsible for bridging many micelles into a condensate. The micelle-like clusters and condensate were dynamic and liquid-like. In addition, Nanog condensates could induce DNA-DNA attraction mediated by micelle-like structures.

## Introduction

Eukaryotic transcription factors (TFs) regulate gene expression by binding to cis-regulatory regions, especially enhancers, and by recruiting other factors, such as coactivators and RNA polymerases. In mammalian embryonic stem cells (ESCs), a small set of TFs called master TFs, including Nanog, Sox2, and Oct4, are known to regulate the expression of many other genes crucial in controlling the ESC state [1][2][3]. At some active gene loci in ESCs, molecular condensates or clusters are found near super-enhancer regions, where master TFs and coactivators, including mediator complexes, are enriched [4][5][6][7]. The levels of active histone markers H3K27ac and H3K4me1 are also higher in these regions [5]. The transcription of these genes is affected by the concentration of the construct [5]. Based on these observations, a phase-separation model for transcription has been proposed [8]. Transcription factors, coactivators, and RNA polymerases form condensed clusters through fuzzy and multivalent interactions. Many components of clusters contain intrinsically disordered regions, which can form condensates via liquid-liquid phase separation (LLPS) [9]. However, the molecular structures and underlying mechanisms of LLPS remain unclear.

Biomolecular LLPS is involved in several cellular functions. However, it is generally difficult to study the details of the molecular structures in condensates using experimental methods owing to the limitation of the temporal and spatial resolutions. Molecular dynamics (MD) simulations are a complementary approach for elucidating the molecular structure of these protein condensates. Molecular dynamics simulations can provide insights into the atomic interactions of individual residues and their roles in driving LLPS. Standard all-atom MD simulations are not feasible for such long-term behavior in large-scale systems. In such cases, coarse-grained (CG) models can effectively accelerate the simulation speed[10], and were applied to simulate LLPS[11][12][13][14]. In the commonly used residue-level CG model, each protein amino acid residue is represented by a mass particle. Globular domains are often treated with structure-based Go-type terms to maintain a native-like folded conformation[15][16]. In addition, sequence-dependent interactions, termed hydrophobicity scale (HPS) potential[11][17], are used to describe physicochemical interactions, such as electrostatic and hydrophobic interactions between disordered regions. The interaction parameters of amino acids have recently been refined towards the LLPS simulations[11] [12][17][18].

In this study, we focused on Nanog, a master TF, to study the molecular mechanism of protein condensates and their effect on DNA. In mouse ESCs, the cytokine leukemia inhibitory factor induces self-renewal [19][20]. Nanog drives the self-renewal of ESCs independently of leukemia inhibitory factor [21][22]. Nanog consists of three domains: the N-terminal domain (NTD), DNA-binding domain (DBD), and C-terminal domain (CTD) (Fig. 1A). The DBD is a globular domain consisting of three α-helices [23]. The NTD and CTD are intrinsically disordered and essential for transactivation [24]. The NTD is highly charged, similar to many transactivation domains of other TFs[22][21][24]. In contrast, the CTD has only a few charged residues (Fig. 1B). Interestingly, it was revealed that in mouse ESCs, the transactivation ability of Nanog CTD was approximately 7-fold higher than that of NTD [24]. The CTD consists of eight to ten tryptophan repeats (WRs), which are five-residue long and begin with tryptophan (Trp). The WR regions are crucial for the function of Nanog in forming homodimers or oligomers [25][26][27][28]. When Trp residues in WRs are mutated to Ala, Nanog cannot form complexes, and leukemia inhibitory factor-independent self-renewal ability is reduced [25][26][27][28]. However, the molecular mechanism of LLPS formation and the relationship between oligomerization and LLPS remain unclear.

**Figure 1.**
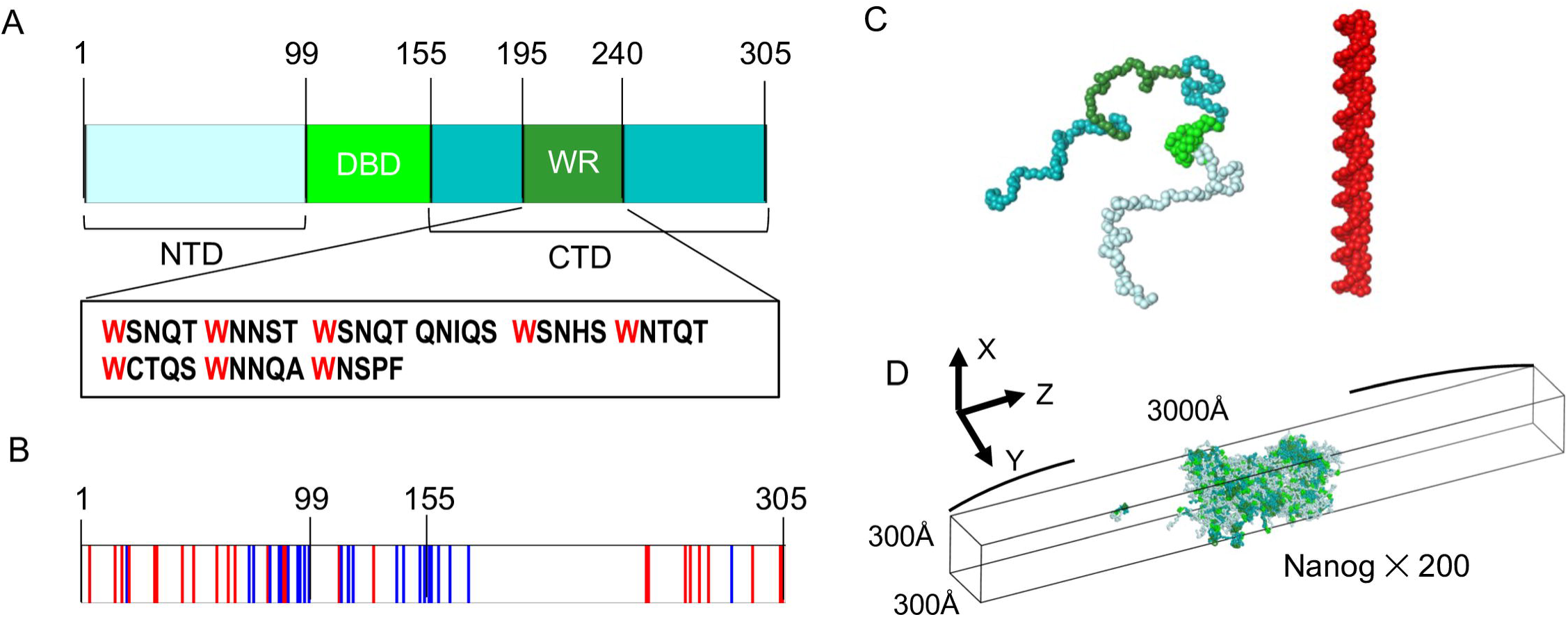
**Nanog structure and simulation setup** (A) Human Nanog sequence. NTD, N-terminal domain; DBD, DNA-binding domain; CTD, C-terminal domain; WR, tryptophan-repeated region. The amino acid sequence in WR is depicted. (B) Charged resides in Nanog. Blue, positively charged residue; red, negatively charged residue. (C) Coarse-grained models of Nanog and DNA. For Nanog, NTD, DBD, CTD, and WR are colored with palecyan, green, teal, and forest, respectively. The same color scheme applies to most images. (D) Simulation setup. In most simulations, 200 Nanog molecules were put in the slab box with the periodic boundary conditions.

In this study, we examined the molecular mechanism of droplet formation by human Nanog using CG MD simulations (Fig. 1C, D). We first verified our model and observed LLPS when the simulation started with condensed and uniformly distributed configurations. We found that Nanog formed condensed clusters primarily via its hydrophobic residues in the CTDs and that the condensate contained micelle-like clusters. These clusters were formed via interactions between WR regions. When we introduced the eight mutations from Trp to Ala, as was done in the previous experiment, the clusters broke, and molecules in the condensate diffused to the dilute phase. The CTD-only mutant formed a more compact condensate than the wild type (WT) did, suggesting that the NTD and DBD are essential for higher fluidity. Finally, we added DNA fragments to the Nanog condensates and observed that the DNA fragments bound to the surface of the clusters. DNA fragments were closer to each other when mediated by Nanog condensates.

## RESULTS AND DISCUSSIONS

### Phase separation of simulated human Nanog

Before investigating the detailed structure and dynamics of human Nanog, we first examined our simulation model to confirm whether it could reproduce the phase separation observed in the experimental results [9]. An *in vitro* experiment showed that Nanog formed droplets in a solution with an average concentration of 10 μM and 10 wt. % polyethylene glycol (PEG) as a crowder [9]. We performed MD simulations for a system containing 200 human Nanog proteins in a slab simulation box of 300Å × 300Å × 3000Å[11], starting from a completely phase-separated configuration (see Methods for the initial configuration setup). We repeated eight independent runs of 5 × 10^7^ MD steps with different stochastic forces.

Figure 2A depicts the time courses of the center of mass of every Nanog molecule along the long axis (z-axis) in one trajectory, indicating that most molecules remain in the condensed phase, and a few molecules leave the condensate (Fig. S1 for the other trajectories, Movie S1). The time courses of the condensed phase size suggest that the trajectories reach near-equilibrium after 2∼4 × 10^6^ MD steps (Fig. S2A, B). The final structure is shown in Fig. 2B. In all the eight trajectories, the phase-separated configurations were stable during the simulations. Once some molecules diffuse into the dilute phase, they diffuse and occasionally merge into the high-density phase. The density of the amino acids along the z-axis in Fig. 2C shows clear evidence of phase separation.

**Figure 2.**
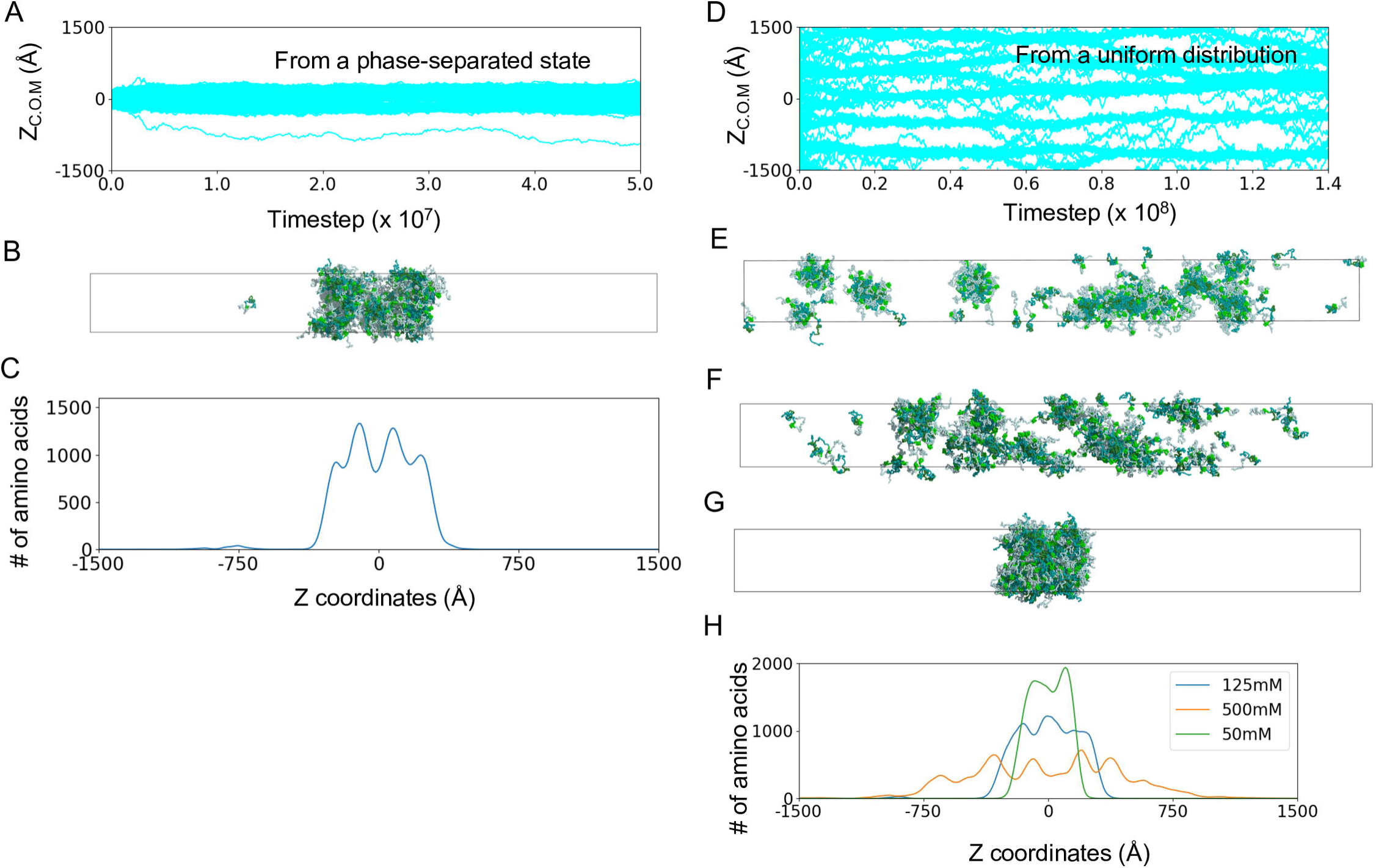
**Model Nanog maintains the phase separated state** (A–C) The results of a representative trajectory starting from a condensed configuration. (A) Time courses of 200 Nanog molecules along the z-axis (long axis) of the slab. Results of other trajectories are shown in Fig. S1. (B) The snapshot at the end of the same simulation as in Fig. 2A. (C) The distribution of Nanog amino acids along the z-axis averaged over 1000 frames in the same trajectory as in Fig. 2A and 2B. (D, E) The results of a representative trajectory started from a random and homogeneous configuration. (D) Time courses of 200 Nanog molecules along the z-axis (long axis) of the slab. (E) The snapshot at the end of the simulation as in Fig. 2D. (F, G) The snapshots at the end of simulations with 500mM (F) and 50mM (G) monovalent ion concentrations. (H) The distributions of Nanog amino acids along the z-axis averaged over 1000 frames in all the trajectories with three monovalent ion concentrations (green: 50mM, blue: 125mM, yellow: 500mM).

The Nanog concentration in the dilute phase of the phase-separated state corresponds to the critical concentration for phase separation. To quantify the concentration in the dilute phase, we counted the number of Nanog molecules in the dilute phase (Fig. S3). In each trajectory, one or two molecules were in the dilute phase at each step, and the average concentration of the dilute phase was 7.64 ± 0.046 μM. This value is consistent with the experimental result that human Nanog forms droplets at an average concentration of 10 μM, thus indicating that the critical concentration for phase separation is below 10 μM [9].

It should be noted that the *in vitro* experiments were performed with 10 wt.% PEG, which can change the phase separation behavior. Therefore, we briefly examined the effect of the crowder, including repulsive spheres, in the simulation box to represent ideal crowders (Fig. S5). The results showed that the condensate was stable during the simulation. Thus, this result does not contradict the experimental results. We note that adding the crowder markedly reduced the Nanog concentration in the dilute phase, thus decreasing the critical concentration for phase separation.

In addition to the above simulation that started from a phase-separated configuration, we also examined another initial configuration in which Nanog molecules were uniformly distributed in the slab simulation box. We performed one MD run with 1.4 × 10^8^ MD steps for this system. Figure 2D shows the time courses of Nanog molecules along the z-axis, which indicates that Nanog quickly formed small clusters during the simulation, which occasionally merged into larger clusters. The final snapshot is shown in Fig. 2E. Clearly, this simulation did not reach equilibrium because of the limitation of simulation time. However, small- to medium-sized clusters were formed, which supports the view that Nanog can form small clusters or condensates, perhaps leading to a phase-separated form.

Previous experimental and theoretical studies demonstrated the effect of salt concentrations and thus roles of electrostatic interactions on LLPS[11][29]. Here, we examined the role of electrostatic interactions to the Nanog droplets by performing MD simulations with a higher and lower salt concentration than the above case of 125 mM. At 500 mM, from a completely-phase separated configuration, we saw gradual dissolution of the Nanog condensate: Many molecules diffused to the dilute phase (Fig. 2F, Fig. S6D and S6E). Similar results were reported for other proteins including Sox2, FUS, and TDP43 in the previous experiments[29]. Interestingly, as shown in Fig. 2F, Nanog molecules did not fall apart to monomers, but remained forming small clusters. The high salt concentration weakened the electrostatic interactions, but did not change the hydrophobic interactions modeled as the HPS potential. Thus, we suggest that, while the condensate is stabilized by electrostatic interactions, small clusters are formed by non-electrostatic interactions. Next, at 50 mM ion concentration, the phase-separated state was stable during the simulations (Fig. 2G, Fig. S6D, S6E). Figure 2G represents the snapshot at the end of one simulation at 50mM, which resembles that at 125mM conditions. However, the maximum density of amino acids was higher at 50 mM than that in 125mM (Fig. 2H). The condensate at 50mM salt was more compact than that at 125mM. The lower salt concentration strengthens the electrostatic interactions that led to higher density condensate.

We conclude that the current simulation model can reproduce the phase-separated form of Nanog observed in an *in vitro* experiment [9]. We suggest that the electrostatic interaction is important to form the Nanog condensate, or droplet, whereas hydrophobic interactions are responsible for forming small clusters.

### Micelle-like clusters in the Nanog condensate

The above observation in our simulations suggests that the condensed structures were not homogenous and were made of many clusters that are bridged to form the condensate. Especially in the simulation starting from a random and uniform distribution, Nanog formed distinctive small and near-spherical complexes (Fig. 2E). Similar structures were observed in the condensate in the simulation of the phase-separated form (Fig. 2B) and in the configurations with higher and lower salt concentrations (Fig. 2F, G). To study the structural features of the condensate, we calculated the inter-molecule residue-residue contact map using a cutoff distance of 6.5 Å (Fig. 3A). As shown in Fig. 3A, we found high frequencies of contact between the hydrophobic regions (155– 240 residues) in the CTD, especially between WR regions. We suspected that a fraction of Nanog molecules in the condensate might form clusters via direct interactions between the WR regions.

**Figure 3.**
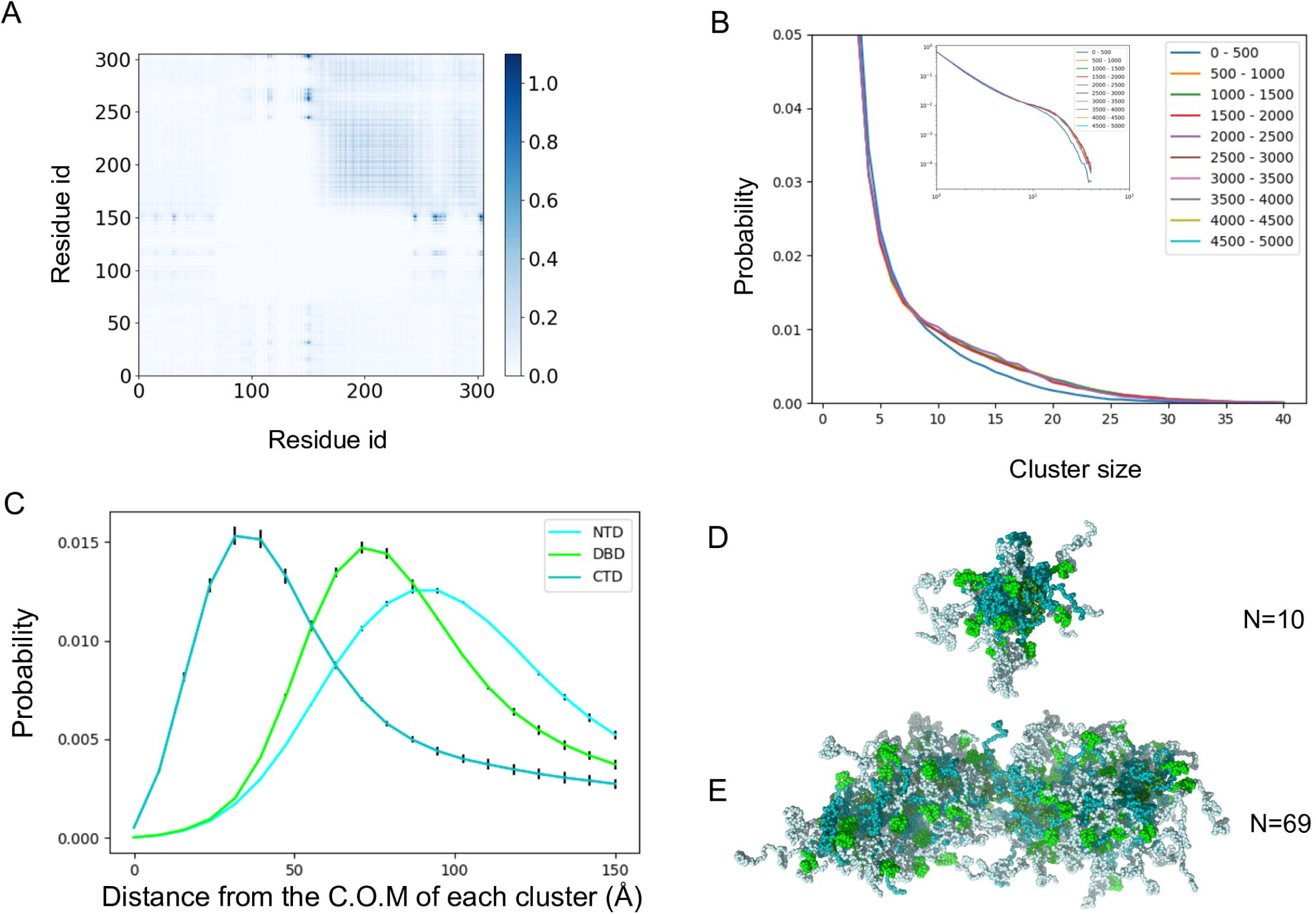
**Nanog forms micelle-like clusters in the condensed phase** (A) The residue contact map between two Nanog molecules. Residue pairs of different molecules within 6.5 Å are considered as the contact. The averaged numbers of contacts in all frames and trajectories are depicted. (B) The distribution of the cluster size in different simulation periods. The entire simulation time is divided into 10 periods. (inset, log-log plot). (C) The distributions of the distances of each domain from the center of mass of the cluster. Cyan, NTD; green, DBD; teal, CTD. (D and E) Snapshot structures of clusters with 10 molecules (D) and the largest cluster containing 69 molecules (E).

To quantify this possibility, we performed cluster analysis based on the contact between WR regions (see MATERIALS AND METHODS for details). The cluster size was distributed broadly from four to 70 molecules (Fig. 3B). The cluster size distribution converged well in the early stages of the simulations (∼5 × 10^6^ MD steps). For each cluster, we calculated the distances of the three domains (NTD, DBD, and CTD) from the center of the cluster (Fig. 3C). The distributions indicate that in the clusters, CTDs are distributed near the center of the cluster, DBDs in the middle, and NTDs near the surface of the cluster. This bias suggests the presence of micelle-like structures. The CTD contains WR regions that attract each other. There were many charged residues in the NTD and DBD, but fewer residues in the CTD. These results are consistent with the typical physicochemical nature of the micelles. Similar to other surfactant molecules[30], we found that when the number of Nanog molecules in a cluster was small, spherical micelles were formed (Fig.3D). Cylindrical micelles were formed when the number of Nanog molecules increased (Fig. 3E).

In the current simulation setup, we included the HPS potential between the disordered regions, but not the DBD, because the HPS potential was developed for the disordered regions. However, we were concerned that this setup might artificially have induced micelle-like structures in our simulations. To test this possibility, two control simulations were performed. First, the HPS potential was applied to all amino acids, asking if this setup also exhibits the micelle-like structure of Nanog in the condensate (Fig. S7). For the second test, we simulated a mutant Nanog that lacked DBD (Note that there is no corresponding experiment at the moment and thus this is merely to test robustness of the micelle-like structure). The results showed micelle-like structures similar to those of the WT (Fig. S8). Therefore, we conclude that the micelle-like structures were not artifacts of the setup.

As mentioned above, we found similar clusters in the configurations with higher and lower salt concentrations. To check if these clusters were micelle-like or not, we did similar analyses for these results (Fig. S6 C, F). In each simulation Nanog formed micelle-like clusters. Interestingly, the micelle-like clusters were stable even in the higher salt concentrations (Fig. S6F). Comparing the contact maps of the medium (125mM) and higher (500mM) salt concentrations, we found that the contacts between NTDs and DBDs were lost in the higher salt concentrations. This result is expected because the NTD has some negatively charged residues and the DBD has many positively charged residues (Fig. 1B).

In summary, we suggest that Nanog forms droplets by two different interactions. First, Nanog forms micelle-like clusters primarily by the hydrophobic interactions between its WR regions. Next, clusters bind each other via electrostatic interactions between the NTD and the DBD.

### Fluidity in the Nanog condensate

To study dynamics of the Nanog condensates, we computed the mean square deviation of each Nanog molecule as a function of the time difference in the condensate as well as in the dilute phase as a control (Fig. S4). Each mean square deviation, as a function of the time difference, was well fitted by a linear line for both the dilute and condensed phases, indicating the normal diffusion. Thus, both the dilute and condensed phases can be regarded as fluids. In our simulations, the diffusion coefficient in the condensed phase was approximately 1/10 of that in the dilute phase.

Next, we tracked selected Nanog molecules in the condensate to examine protein fluidity in micelle-like clusters as well as in the condensate (Fig. 4). Figure 4A and 4C represent movements of Nanog molecules that were in a selected region along z-axis at a time point 2.0 × 10^7^, as a proxy to the fluorescence recover after photobreaching (FRAP) experiment. Note that all the selected molecules were not in one micelle. The molecules diffused nearly everywhere within the condensate in approximately 10^7^ MD steps. Next, Fig. 4B and 4D represent tracking of 11 Nanog molecules involved in a micelle-like cluster at a time point of 2.3 × 10^7^ MD steps. We observed a dynamic rearrangement within the condensed phase. The number of Nanog molecules remained in the same cluster slowly decreased; six and three molecules after 0.4 × 10^7^ and 0.8 × 10^7^ MD steps, respectively from the reference time point. Within the cluster, the relative diffusion was restrained, by definition. Once ejected from the cluster, molecules started diffusing freely from the cluster.

**Figure 4.**
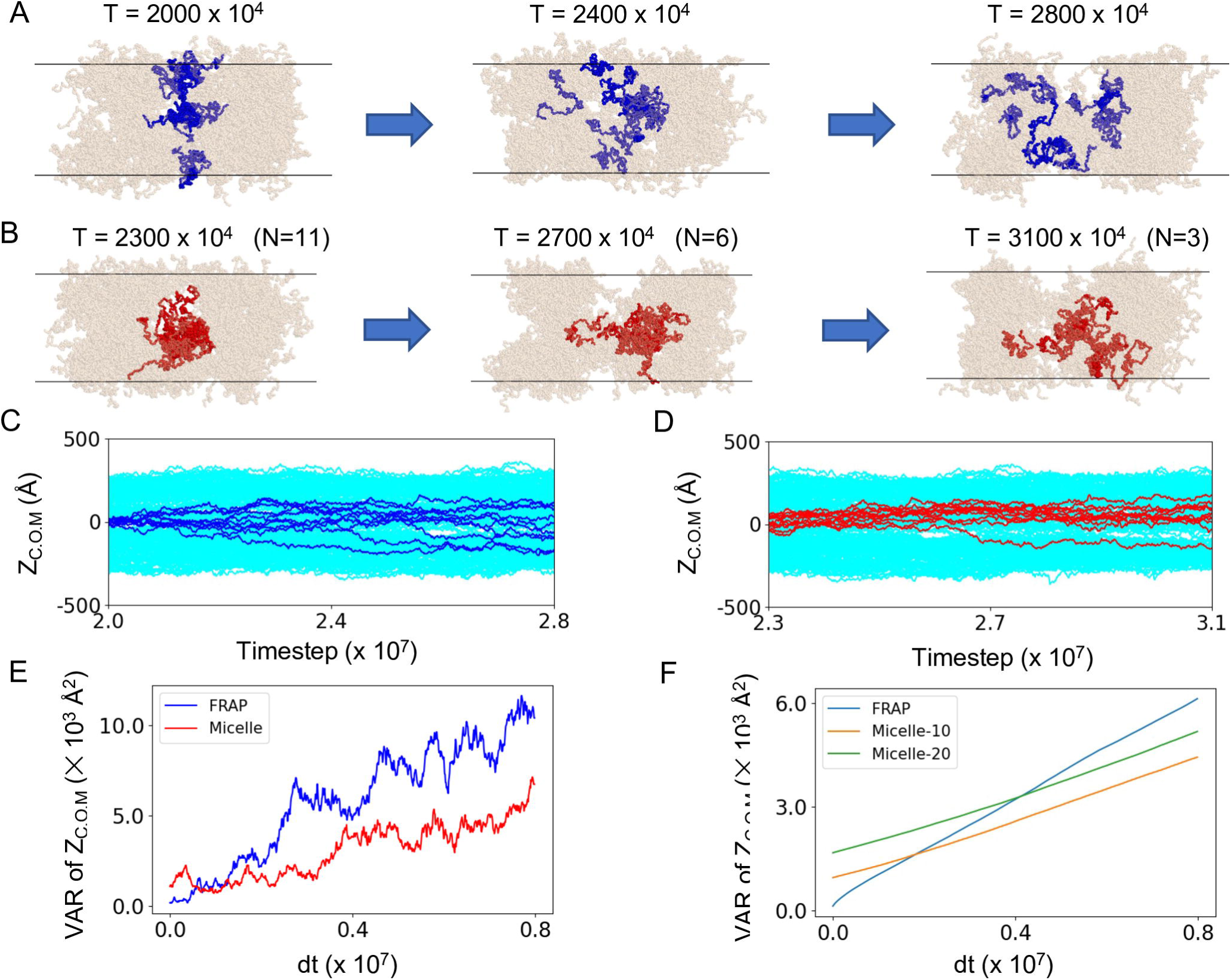
**Micelle-like clusters are dynamic and fluid** (A and B) Three successive snapshots in a simulation period with tracked molecules indicated in blue and red. (A) Molecules near the center of the condensed phase at the 2.0 × 10^7^ step are marked in blue (left) and tracked in the future direction (FRAP-like plot). (B) Molecules in one micelle-like cluster at the 2.3 × 10^7^ step are marked in red and are tracked in the downstream analysis. N in the parentheses represent the number of Nanog molecules remaining in the same cluster as the initial time point. (C,D) Time courses of the assigned blue (C) and red (D) molecules along the z-axis of the slab. (E) Variance of the z-coordinates of the center of mass of tracking molecules as a function of the time duration from the reference time point. Blue; those in A and C, red; those in B and D. (F) Averaged variance of the z-coordinates as a function the time duration from the reference time point. For the sample from one micelle, we compared the spreading of a small micelle containing 9 to 11 molecules (orange) with that of a large micelle containing 20-30 molecules (green).

To compare the diffusive spreading from one micelle and from one region not restricted to the micelle for the same pair of trajectories, we plotted in Fig. 4E the variance of the z-coordinates in the tracking molecules as a function of the time duration from the reference time points. Clearly, the variance increased faster for the sample from one region (termed FRAP) than that from one micelle (denoted as micelle). For the latter, the variance stayed nearly constant for a short period at the beginning, ∼0.3× 10^7^ MD step, during which all the molecules remained in the same micelle. Next, we obtained the averaged variance as a function of the time duration over all the trajectories in Fig. 4F. We found that the diffusive spreading from one micelle is clearly slower than that from one region (FRAP). The spreading rate is not markedly different for different sizes of micelles; micelles containing 9-11 molecules and those containing 20-30 molecules showed very similar spreading rates. In summary, while we observed translocations of Nanog molecules from one cluster to another, the inter-cluster transition is markedly slow, relative to normal diffusion in the condensate.

In summary, micelle-like structures are not fixed, but dynamically change their component molecules. We suggest that although the condensates are not uniform and have micelle-like substructures, these underlying substructures are transient and dynamic. Within a micelle-like condensate, the molecular motion can differ from normal diffusion. Molecules involved in one micelle can be restricted to the micelle-like structures on a short time scale. Although molecules can depart from one micelle to reach another in the long term, it would involve an escape process with some free energy barrier.

### Tryptophan in the WR region is essential for the LLPS

The importance of Trp residues in the WR region has been demonstrated [27][28]. To examine the effect of Trp residues in the WR region on LLPS ability, we constructed a mutant model in which all eight Trp residues in the WR region were mutated to Ala (W8A) and performed the same simulation as above, starting from a totally phase-separated configuration. As shown in Figure 5A, the condensed phase dissolved quickly during the simulation of the W8A system. The W8A molecules diffused throughout the slab simulation box (Fig. 5B, C). We did not find any stable clusters or oligomers. Although the simulation was not long enough to reach a completely uniform state, a sufficiently long simulation would lead to a uniform distribution in the simulation box. The simulation clearly showed that Trp residues in the WR regions are essential for the formation of LLPS.

**Figure 5.**
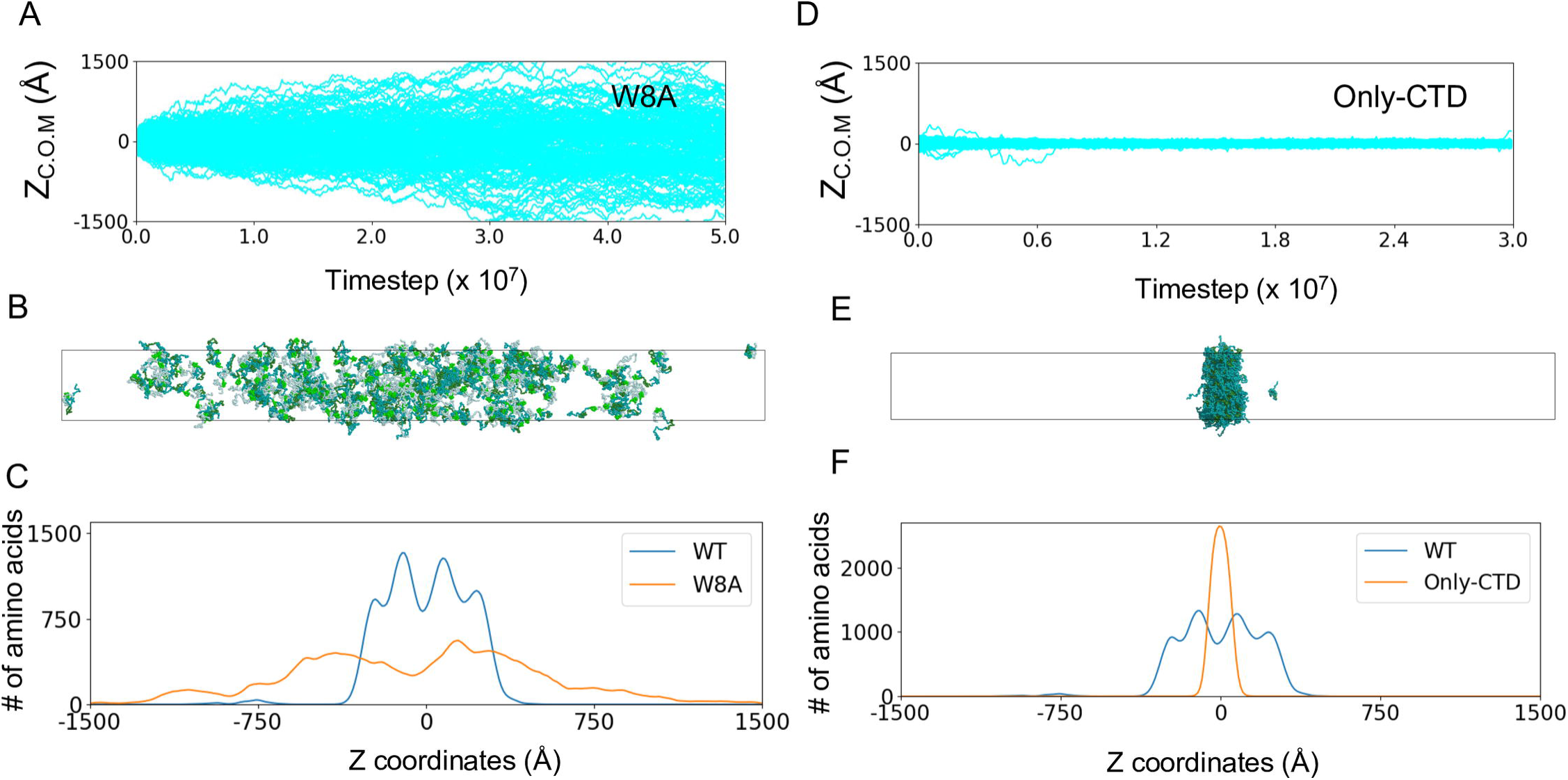
**Mutant Nanog simulations** (A–C) Simulation results for the Nanog mutant W8A. (A) Time course of 200 Nanog molecules along the z-axis (long axis) of the slab. (B) Snapshot at the end of the simulation. (C) Density of amino acids along the z-axis. Yellow, W8A; blue, WT. (D–F) Simulation results of the Nanog deletion mutant Only-CTD. (D) Time courses of 200 Only-CTD Nanog molecules along the z-axis (long axis) of the slab. (E) Snapshots taken at the end of the simulation. (F) Density of amino acids along the z axis. Yellow, Only-CTD; blue, WT.

### Phosphorylation of Nanog slightly weakens the condensate

Nanog contains several serine residues that are known to be phosphorylated in human ESCs[31][32]. While the phosphorylation at the residues in the DBDs were well-studied, the effect of the phosphorylation in the NTDs remains unclear[32]. Previous studies about FUS showed that the phosphorylation destabilized the droplets[33]. To examine the effect of phosphorylation in the N-terminal domain on the droplet we constructed the mutant that changed the charge of Ser52, Ser56, Ser57, and Ser65 to -1.0 (Fig. S9A). These serine residues are well conserved and known to be phosphorylated[34][31]. The condensate of the phosphorylated Nanog was stable during the simulations (Fig. S9B, C). We found that the condensed configurations in the final frame have slightly more cavities than that of the unphosphorylated Nanog, and the density of amino acids also shows similar extension of the condensates (Fig. S9D, E). As mentioned above, the NTD has some negatively charged residues and the interaction between the NTD and DBD was important for the formation of droplets. The net charge of human Nanog in its unphosphorylated form is -1, and thus the phosphorylation changes the net charge to more negative (-5 in the current setup). We found that the phosphorylation decreased the contacts between the NTD and DBD (Fig. S9F). The phosphorylation in the NTD changes the charge balance between the NTD and the DBD and strengthens the repulsion between the NTDs. Previous studies showed that the mutation of the phosphorylation sites upregulated the reprogramming function of Nanog[31]. We suggest that the phosphorylation in the N-terminal domain destabilizes the droplets of Nanog, which should reduce the transcription of the target genes. We note that, given that the pKa value of the secondary ionization of the phosphate group is close to 7, the phosphate group can have -1.0 or -2.0 charges depending on its local environment. In our simulation, we used -1.0 to represent modest effects of phosphorylation. With the charge of -2 in the phosphorylated serine, we would expect even stronger effects than the current case.

### NTD and DBD form the cavity in the condensed configuration

The NTD and DBD were located on the surface of the micelle-like clusters of Nanog. These surface-exposed regions contain many hydrophilic and charged residues. This implies that repulsive forces can exist between the clusters. We presumed that these repulsive forces might create void spaces in the condensates, thereby enhancing fluidity. To test these propositions, we constructed a mutant that lacked the NTD and DBD (Only-CTD), and performed the same simulation as above. During the simulations of this mutant, the condensate was stable, as was the case for the WT-Nanog (Fig. 5D). However, the internal structure was distinct from that of WT-Nanog; As shown in Figure 5E, which is a snapshot of the final frame, we found a highly compact condensate without any significant cavity. Quantitatively, we calculated the density of amino acids along the long axis of the slab (Fig. 5F) and found that the density of amino acids in Only-CTD was much higher than that of amino acids in WT-Nanog. By deleting the NTD and DBD, which contain many hydrophilic and charged residues, the mutant became highly compact. In the case of WT-Nanog, small micelle-like clusters packed loosely to form a large condensate because of the relatively hydrophilic and charged NTD and DBD.

In a previous study, the CTD of Nanog alone was insoluble in water[28]. Our results indicated that WT-Nanog could dissolve in water owing to the formation of micelle-like structures.

### Condensates induce DNA-DNA attraction

Transcription factors form droplets along with chromatin, transcription machinery, and coactivators in a model of transcription condensates. To study the effects of the Nanog LLPS condensates on DNA, we added four 50 bp DNA fragments of the poly CG sequence to the neighbors of the condensate that contained ∼200 Nanog molecules. We note that the Nanog consensus sequence is far from the poly CG sequence. Here we used the poly CG sequence only to examine non-specific binding of Nanog via electrostatic interactions. During the simulation, the DNA fragments were incorporated into the Nanog condensate (Fig. 6A). Figure 6B, a snapshot of the final time, shows that DNAs are not located in the center of the condensate but are bound to the surface of the condensate. Figure 6C shows a close-up view of the micelle-like cluster in which one DNA fragment is bound to the surface. In the micelle-like clusters of Nanog, DBDs are exposed on the surface of the clusters so that DNA segments can access the DNA-binding domains of many Nanog molecules. We suggest that the micelle-like structure makes a DNA segment to easily bind to multiple Nanog molecules and to co-stabilize the complex.

**Figure 6.**
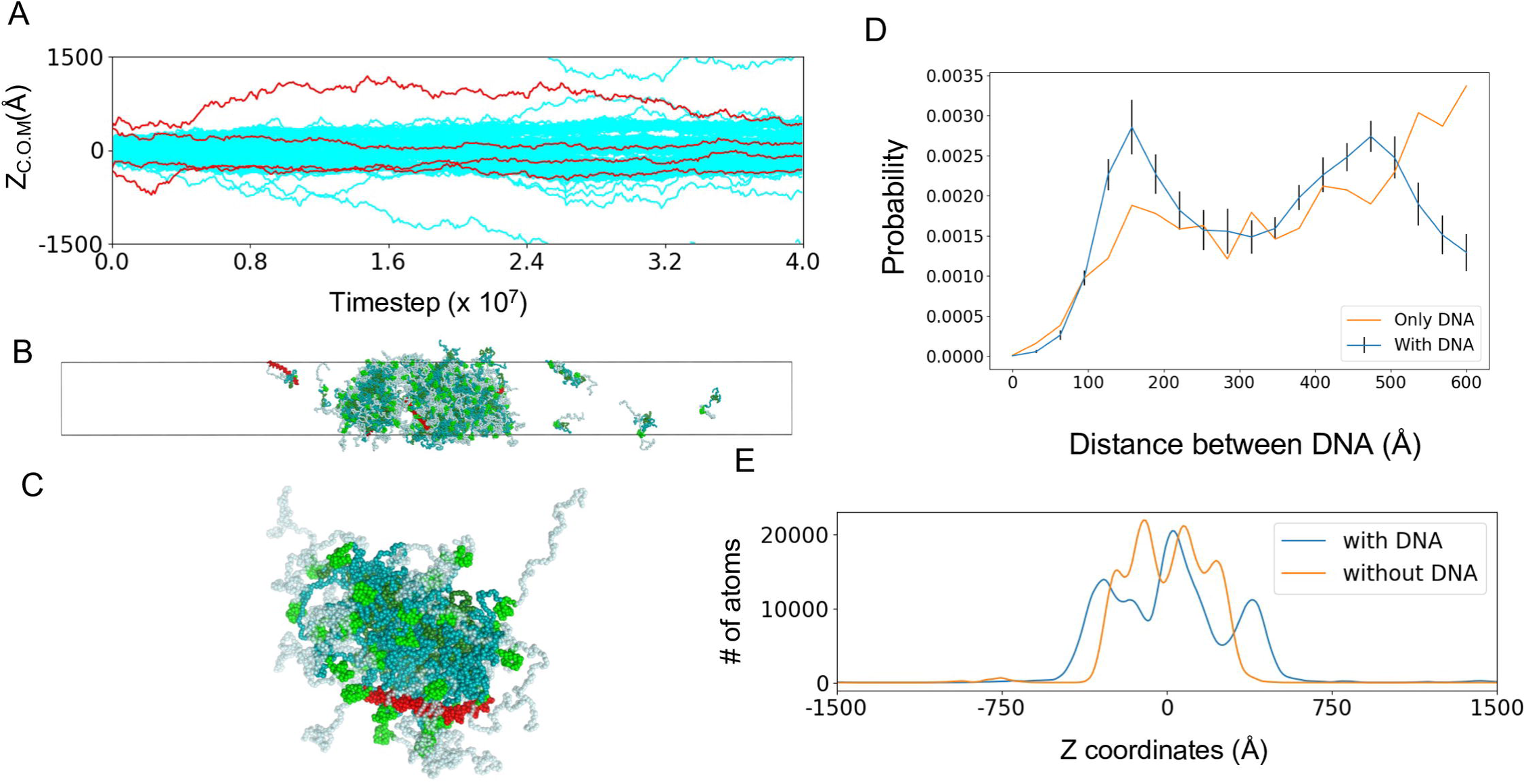
**Condensates containing Nanog and DNA fragments** (A–C) Results of a representative trajectory of 200 Nanog molecules and four DNA segments starting from a fully phase-separated configuration. (A) Time courses of 200 Nanog molecules (cyan) and four DNA (red) along the z-axis (long axis) of the slab. (B) Snapshot at the end of the simulation. Red, DNA (C) Snapshot of a single cluster and a DNA fragment that binds to the cluster. This structure is part of the final configuration. (D) Distributions of the distances between the center of mass of DNA fragments with and without Nanog condensates. Blue, case with Nanog condensate; yellow, case without Nanog. (E) Distribution of atoms calculated from each type of coarse-grained particles including Nanog and DNA along the z-axis in the simulations of 200 Nanog with and without DNA.

In the simulations, multiple DNA segments were bound to the Nanog condensate, making the distance between the two DNA segments relatively close (Fig. 6A). To quantify this, we compared the DNA-DNA distance distribution in the Nanog condensate with that in a DNA-only system, where only four 50 bp DNA segments were simulated in the same slab simulation box (Fig. 6D). With the Nanog condensate, we found a prevalence of DNA-DNA centroid distances in the range of 100–200 Å. However, we did not observe this in the DNA-only system. In the micelle-like structure, the DBDs tended to be located at ∼80 Å from the center of the micelles (Fig. 3C). Thus, two DNA segments bound to the opposite end of the spherical micelle-like cluster had a distance of ∼160 Å. Micelle-like clusters can bind multiple DNA segments at once; therefore, the cluster makes the DNA fragments closer than in the DNA-only system. This is in agreement with recent experiments and with the transcriptional condensate model, which states that transcription factor condensates bring super-enhancers and promoters into proximity [35]. The DNA-DNA distance distribution has the second peak near 450 Å, which corresponds to the breadth of the Nanog condensate in the z-direction of the slab. DNA weakly favored the interface of the condensate, which resulted in the peak of the distribution corresponding to the breadth of the condensate.

We compared the density of atoms calculated from each type of the CG particles containing Nanog and DNA along the z-axis for simulations with and without DNA as shown in Figure 6E. The Nanog condensate containing DNA was slightly more extended than that without the DNA. Some Nanog DBDs that are bound to DNA cannot have the same attractive interactions to Nanog NTDs as in the case without DNA, which weakens the interactions between micelle-like clusters. The net negative charges of the condensate may also weaken the condensate.

To examine the saturation effect of DNA fragments, we added four more DNA fragments to the final configurations of the above simulation and extended MD simulations for 5.0 × 10^7^ MD steps. During the simulations few DNA fragments entered the droplets; two of the four added DNA were completely dissociated, the other two resided near the interface of loose condensate (Fig. S10A, B). The snapshot at the end of the simulation showed large cavities in the droplet (Fig. S10C). The net charge of the condensate was highly negatively charged, and thus have overall electrostatic repulsions, which tends to break the droplet. The atomic densities along z-axis showed no significant difference between the cases containing four and eight DNA fragments (Fig. S10D). A previous experiment showed that a condensate of positively charged proteins is enhanced with a low density of RNA, but is dissolved with a high-density of RNA[36]. The current result may qualitatively correspond to the high-density case since Nanog has the net charge of -1.

### Biological significance

Nanog is one of the master transcription factors and regulates many genes cooperatively with other transcription factors such as Oct4 or Sox2 in mammalian ESC. Recent experiments showed that human Nanog forms condensate, or droplet, at the concentration 20μM[9]. Below the critical concentration to form the condensate, previous experiments identified formation of dimer as well as broad range of oligomers[26]. Our simulation study characterized structural feature in the Nanog condensate; human Nanog forms ∼100Å scale micelle-like structures, which are weakly bridged together to form larger-scale condensate. Hydrophobic interactions of the WR region, especially tryptophan, in the C-terminal domain are responsible for the micelle formation, whereas assembly of micelles into condensate is driven by electrostatic interactions in the N-terminal and DNA-binding domains. This micelle-like structure can have advantages for Nanog functions and explain some results obtained in previous studies.

First, we discuss possible states of Nanog in the cell. We estimated the critical concentration for the condensate formation as 7.6μM. While this is a rough estimate given the simplicity of the simulated model and the small number, i.e., 200, of Nanog molecules in the simulation system, we guess the real critical concentration would be between 1 and 10μM. Although the concentration of Nanog in ESC has not been characterized to our knowledge, the concentrations of other master TFs, Sox2 and Oct4, are known to be on the order of 1μM[37][38][39]. Therefore, Nanog alone unlikely to form the condensate in ESC. Yet, given the dimer or oligomer formation reported previously, we consider that the micelle-like cluster remains stable somewhat under the critical concentration for the condensate formation. Our simulation started from a random and uniformly distributed configuration showed prompt formation of micelle-like clusters of Nanog in the early stages (Fig. 2D), which would support formation of clusters even at lower concentration than the critical one. We are not sure if Nanog alone forms clusters or not in the ESC. Importantly, the critical concentration for the Nanog condensate is only modestly higher than the Nanog concentration in ESC. Enhancer regions of Nanog-target genes have multiple Nanog-binding sites in the super-enhancers, which would induce formation of either cluster or condensate.

Furthermore, the critical concentration should be further modulated by chemical modification such as phosphorylation. Our quick examination suggested that the phosphorylation of Nanog NTD sites slightly increases the critical concentration of the condensate formation. Thus, the phosphorylation would reduce the condensate formation for a given concentration and thus would down-regulate the target transcription. This result is, at least qualitatively, consistent with the experiment which showed that the phosphorylation-deletion mutant enhanced the Nanog activity[31]. Notably, our simulation showed that the cluster formation mediated by the CTD domain is not affected by the phosphorylation in the NTD.

The micelle-like structure can be efficient for Nanog function. In our model the CTDs interact with each other by hydrophobic interactions, and NTDs and DBDs are exposed to the surface of the cluster. Thus, DNA can easily be accessible to the DBD of Nanog. In addition, one DNA segments can bind multiple Nanog molecules, at least via sequence non-specific electrostatic interactions. The Nanog-DNA complex in the Nanog cluster can bring different DNA regions closer as shown in Fig. 6C and 6D. Previous studies using fluorescence resonance energy transfer experiments showed that Nanog complex could bring DNA fragments closer and the ability was lost by the mutation of tryptophan residues to alanine[28]. Nanog micelle-like cluster can bring enhancer and promoter regions closer in space.

Previous studies revealed that the mutant containing only CTDs is insoluble in aqueous solution[28]. As shown in Fig. 5E, the mutants containing only CTDs formed markedly more compact structures than the full-length Nanog. We suggest that the charged NTDs and DBDs exposed to the solution in the micelle-like structure prevents CTDs from forming insoluble aggregates.

A recent study proposed that the CTDs of Nanog can form cross-β structures with the same domains in different Nanog molecules and finally form fiber-like structures[28]. In the micelle-like structures, CTDs of several molecules are assembled at the center with high density. We speculate that within such a high-density mixture of CTDs, the WR region could form cross-β structure after some latent time. Our simulation that used residue-revel coarse-grained models cannot faithfully represent hydrogen-bonds interactions, making simulation of β-sheet formations intractable. Fully atomistic models are required to examine the roles of cross-β structures, which is left for future study.

## CONCLUSIONS

In this study, we investigated the molecular structures of condensates formed by human Nanog using CG MD simulations. We first verified the model and observed the LLPS when the simulation started from condensed and uniformly distributed configurations. We observed that Nanog formed condensates primarily via hydrophobic residues in the CTDs. We found that the condensate contained micelle-like clusters formed via interactions between the WR regions. When we added DNA fragments to the Nanog condensates, we found that the DNA fragments bound to the surfaces of the clusters could become closer to each other. At the same time, the Nanog condensate became less packed in the system with DNA.

The current study focused on the protein condensate formed by Nanog alone; however, chromatin should contain many types of TFs, coactivators, RNA, and RNA polymerases. How mixtures of TFs and other elements alter the nature of condensates is of interest for future studies.

## MATERIALS AND METHODS

### Molecules

We used full-length human Nanog protein, which contains 305 residues. For double-stranded DNA, we used a 50 bp poly-CG sequence.

### Coarse-Grained Model for Nanog and DNA

We used residue-level CG models for large-scale MD simulations of Nanog condensates, with and without DNA [40]. For Nanog, each amino acid was represented by one bead located at the Cα atom position, whereas each nucleotide in the DNA was represented by three beads, each representing a phosphate, sugar, and base.

The potential energy functions for Nanog consist of the AICG2+ potential [41], HPS potential[11][17], excluded volume interactions, and Debye–Huckel electrostatic interactions. For the DBD of Nanog, we used structure-based potentials from the AICG2+ model to maintain the folded structure [42]. For the disordered regions of Nanog, we used the flexible-local potential. For nonlocal interactions between disordered regions of both intra- and intermolecular interactions, we used the HPS potential. The HPS model has been used previously with progressively refined parameters for the simulations of LLPS of disordered proteins [43][44][45][29]. In the present study, we selected the parameter set obtained by Tesei et al. [17].

For DNA, we used the 3SPN.2C model developed by de Pablo et al. [46], which has been used together with the AICG2+ model in previous studies [47][48][49]. The excluded volume and Debye–Huckel electrostatic interactions were considered for Nanog-DNA interactions.

### Molecular Dynamics Simulation

As in previous simulation studies of LLPS, we used a slab simulation box (small x and y side lengths and a large z side length) with the periodic boundary condition. The slab box is superior to the more standard cubic box in that the slab box reduces the finite-size artifact. With a standard cubic box, the droplet, once formed, has spherical surface. For limited size of systems, significant fraction of molecules is involved to the interface of the two phases. Since the interface molecules have frustrated interactions, they lead to energy cost, or the surface tension. This gives rise large finite-size effect. On the other hand, with small x and y side lengths, molecules can form condensate spanning all x and y ranges. With the periodic boundary condition, the interface disappears in x and y directions. To represent the inhomogeneous two phases, we do need long z side length with the interface. The area of one interface is only the product of the side lengths of x and y, which is relatively small in the slab box.

We used a slab box with x, y, and z side lengths of 300, 300, and 3000 Å, respectively. From a preparatory simulation of one Nanog molecule, we estimated the largest residue-residue distance within one Nanog molecular was around 150Å. Thus, we used 300 Å as the x and y side lengths to prevent the molecules from interacting with their periodic images. As for the z side length, too large side length makes reaching equilibrium intractable. We checked the Nanog single molecule can diffuse ∼ 3000 Å within the possible simulation time. In the slab box, we included 200 molecules of human Nanog, with or without four DNA fragments, which corresponds to an average Nanog concentration of 1.23 mM. This average concentration is in between those of the low- and high-density phases of Nanog, which ensures the phase separation.

We used the MD simulation software GENESIS, which contains all models described above[50][51] [52]. All simulations in this study used Langevin dynamics at a temperature of 300 K. We used the GENESIS-CG-TOOL software to create the initial DNA structures [52].

In most simulations, we used completely phase-separated configurations as starting configurations for efficient sampling. To create the configurations, we first performed short MD simulations for the system that contained one Nanog molecule and took an ensemble of conformations. We chose highly compact structures from the ensemble and placed 200 copies into a sufficiently large simulation box with the same length in the x and y axes and with no overlap between amino acids. Subsequently, the box size was gradually decreased by 0.1 Å every 100 MD steps along the z axis to the size which was small enough for all molecules to be in the condensed phase. In the shrinking simulation we used the same force field as used in the sampling and decreased the box size slowly, so that interactions are locally well relaxed. For the simulation, starting from a homogeneous and random configuration, we placed 200 copies in the simulation box to avoid molecule overlap.

As product MD simulations, we performed eight, two, and two independent MD runs for monovalent salt concentrations of 125, 50, and 500mM, respectively, for WT Nanog from a completely-phase separated configuration. The system that contains four DNA fragments was conducted five times, whereas that containing eight DNA fragments was run two times. For the simulation with phosphorylated Nanog, we conducted two MD runs. For the rest of simulations, such as that from a random configuration of WT and the W8A mutant, we performed one MD run each.

### Analysis of Simulation Results

To calculate the concentration in the dilute phase, we performed clustering based on the distance between molecules. In each frame, we first defined the distance between the molecules by the minimum distance between the amino acids in the molecules and calculated the contact map. We assumed that there were no interaction energies between the molecules in the dilute and condensed phases; therefore, we defined the cutoff distance of the contact map as the cutoff distance of the HPS model (50 Å). Based on the clustering results, a molecule in the dilute phase was defined as one that was not in the clusters of five or more molecules.

We used a similar method to study the micelle-like clusters formed by interactions between WR regions. In this case, the cutoff distance to define the contact between the WR regions was 6.5 Å.

In most analysis, we used the Python library MDAnalysis to read and analyze the DCD files[53][54].

### Effect of Crowders

In a previous experiment, PEG8000 was used as a crowder to increase the concentration of the macromolecules. To mimic the effect of the crowder, we performed simulations in the system that contained crowders at a concentration equal to the experimental conditions and compared it with the experimental results. The concentration of PEG8000 in the experiment was 10% (weight percent) [9], corresponding to 2030 repulsive spheres in our simulation box.

To briefly examine the effect of PEG8000, we employed a simple model of PEG8000, with a sphere of radius *R* =19 Å, estimated from the radius of gyration *Rg* of PEG8000 ∼15 Å [55] as 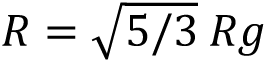 [56]. Repulsive spheres representing PEG8000 were included in the solution as crowders. The interactions between the PEG spheres and proteins were the only excluded volume terms.

We performed simulations under these conditions and found that Nanog formed a condensate with similar fluidity and residue-level contacts to the case without the crowders (Fig. S3D). We suggest that the crowders have minor effects on the structure of the phase-separated Nanog condensate. In contrast, the Nanog concentration in the dilute phase was markedly reduced by the inclusion of the crowders (Fig. S3B). However, the repulsive sphere is an ideal crowder model, which is too simplistic for the PEG model [57]. A more realistic modeling of the PEG in our system is left for future work.

## Acknowledgements

The authors thank Soundhararajan Gopi and Giovanni B Brandani for helpful discussions. This research used computational resources of supercomputer Fugaku provided by the RIKEN Center for Computational Science through the HPCI System Research project (Project ID: JPMXP1020200101). The computer resources are provided by the HPCI system research project (project ID: hp200135, hp210177, and hp220170).

## Author Contributions

### Conceptualization

Azuki Mizutani, Shoji Takada

### Data Curation

Azuki Mizutani

### Formal Analysis

Azuki Mizutani

### Funding Acquisition

Yuji Sugita, Shoji Takada

### Investigation

Azuki Mizutani

### Methodology

Azuki Mizutani, Cheng Tan

### Project Administration

Shoji Takada

### Resources

Yuji Sugita, Shoji Takada.

### Software

Azuki Mizutani, Cheng Tan

### Supervision

Shoji Takada.

### Validation

Azuki Mizutani

### Visualization

Azuki Mizutani

### Writing – Original Draft preparation

Azuki Mizutani

### Writing – Review & Editing

Cheng Tan, Yuji Sugita, Shoji Takada

## Supporting information captions

### S1. Appendix. The detail about the analysis

**S1. Figure. Other trajectories of WT human Nanog simulations**

(A∼G) The results of the MD simulation for 200 Nanog with the same setup as Fig. 2 but with different random numbers. Cyan lines in each figure represent the trajectory of z-axis of centroids for each molecule during the simulation.

**S2. Figure. Estimate of the convergence**

(A)The time course of the size of the condensed phase. (B)The time course of the number of molecules in the condensed phase. In (A) and (B), the blue curve represents the average of all the eight trajectories, and the cyan represents the standard error.

**S3. Figure. Estimate of the Nanog concentration in the dilute phase**

The distribution of the concentration of molecules in the dilute phase. The horizontal axis represents the concentration of the molecules in the dilute phase, calculated from the volume of the dilute phase and the number of molecules in the dilute phase. The distribution was calculated for each trajectory, which was then averaged. Each error bar represents the standard deviation in eight trajectories. The vertical dashed line represents the experimental upper limit of the critical concentration (10μM)[9].

**S4. Figure. The MSD of Nanog molecules**

The mean square deviation (MSD) of a Nanog molecule as a function of the time difference. Comparison of the simulation results for a single Nanog molecule (orange) and the condensed state (cyan). Filled regions represent the standard deviation in all the eight trajectories. The dashed lines are obtained by the liner regression.

**S5. Figure. Simulations with a PEG model**

(A) The model of human Nanog and the PEG8000. The model of Nanog is the same as that in Fig. 1C. NTD, DBD, CTD, WR are colored with palecyan, green, teal, forest. The gray sphere represents a model for one molecule of PEG8000. (B) The result of one trajectory. Cyan curves represent the trajectories of z-axis of the centroids of 200 Nanog molecules. (C) The snapshot at the last frame of the same trajectory. We drew PEG8000 spheres with the radius smaller than the actual one. (D) The inter-molecule contact map. We used the cutoff distance 6.5Å between the two residues. The color represents the average number of contacts over all the frames and all trajectories. (E) The MSD as a function of the time difference. The yellow line represents the MSD of Nanog in the condensed phase with PEG. The blue line is the same result as in Fig. S4.

**S6. Figure. Simulations of 200 Nanog in the slab box with 50mM NaCl or 500mM monovalent ion concentrations**

(A, B) Time courses of 200 Nanog molecules along the z-axis (long axis of the slab) in the simulations with 50 mM condition. Figures A and B are results of the same setup with different random number seeds. (C) The residue contact map between two Nanog molecules in the simulation with 50 mM conditions. (D, E) Time courses of 200 Nanog molecules along the z-axis (long axis of the slab) in the simulations with 500 mM condition. Figures C and D are results of the same setup with different random number seeds. (F) The residue contact map between two Nanog molecules in the simulation with 500 mM NaCl conditions. Residue pairs of different molecules within 6.5Å are considered as the contact in (C) and (F).

**S7. Figure. Simulations of 200 Nanog in the slab box in which the HPS potential was applied to all the amino acids including DBD**

(A) Time courses of 200 Nanog molecules along the z-axis (long axis) of the slab. (B) The snapshot at the end of the same simulation as in A. (C) The residue contact map between two Nanog molecules. Residue pairs of different molecules within 6.5 Å are considered as the contact. The averaged numbers of contacts in all frames and trajectories are depicted. (D) The distributions of the distances of each domain from the center of mass of the cluster. Cyan, NTD; green, DBD; teal, CTD.

**S8. Figure. Simulations of 200 Nanog mutant that lacks DNA binding domain**

(A) Time courses of 200 mutant molecules along the z-axis of the slab. (B) The snapshot at the end of the same simulation in A. (C) The residue contact map between two Nanog molecules. Residue pairs of different molecules within 6.5 Å are considered as the contact. The averaged numbers of contacts in all frames and trajectories are depicted.

**S9. Figure. Simulations of 200 Nanog phosphorylated at four serine residues in the N-terminal domains.** (A)The mutated residues in the N-terminal domain of Nanog. The red colored serine were phosphorylated to have the charges -1.0. (B, C) Time courses of 200 Nanog mutants along the Z-axis (long axis) of the slab. Figure B and C represent results of the same setup with different random number seeds. (D) Snapshot of the end of the same simulations as shown in Figure A. (E) Density of amino acids along the z-axis. yellow, phosphorylated; blue, WT. (F) The residue contact map between two phosphorylated Nanog molecules in the simulation. Residue pairs of different molecules within 6.5Å are considered as the contact.

**S10. Figure. Simulation of 200 Nanog with eight poly-CG DNA fragments.**

(A, B) Time courses of 200 Nanog molecules (cyan) and eight DNA (red) along the z-axis (long axis) of the slab. (C) Snapshot at the end of the same simulation as (A). (D) Atomic density calculated from each type of CG particles including Nanog and DNA along the z-axis in the simulations of 200 Nanog without DNA and with four or eight DNA fragments. (Blue, with four DNA; Green, with eight DNA; Orange, without DNA)

**S1. Movie. Trajectory of the simulation of 200 molecules of human Nanog in the slab box.**

## References

1. Young RA. Control of the embryonic stem cell state. Cell. 2011;144: 940‒954. doi:10.1016/j.cell.2011.01.032

2. Ng HH, Surani MA. The transcriptional and signalling networks of pluripotency. Nat Cell Biol. 2011;13: 490‒496. doi:10.1038/ncb0511-490

3. Orkin SH, Hochedlinger K. Chromatin connections to pluripotency and cellular reprogramming. Cell. 2011;145: 835‒850. doi:10.1016/j.cell.2011.05.019

4. Kagey MH, Newman JJ, Bilodeau S, Zhan Y, Orlando DA, Van Berkum NL, et al. Mediator and cohesin connect gene expression and chromatin architecture. Nature. 2010;467: 430‒435. doi:10.1038/nature09380

5. Whyte WA, Orlando DA, Hnisz D, Abraham BJ, Lin CY, Kagey MH, et al. Master transcription factors and mediator establish super-enhancers at key cell identity genes. Cell. 2013;153: 307‒319. doi:10.1016/j.cell.2013.03.035

6. Li J, Dong A, Saydaminova K, Chang H, Wang G, Ochiai H, et al. Single-Molecule Nanoscopy Elucidates RNA Polymerase II Transcription at Single Genes in Live Cells. Cell. 2019;178: 491–506.e28. doi:10.1016/j.cell.2019.05.029

7. Li J, Hsu A, Hua Y, Wang G, Cheng L, Ochiai H, et al. Single-gene imaging links genome topology, promoter‒enhancer communication and transcription control. Nat Struct Mol Biol. 2020;27: 1032‒ 1040. doi:10.1038/s41594-020-0493-6

8. Hnisz D, Shrinivas K, Young RA, Chakraborty AK, Sharp PA. A Phase Separation Model for Transcriptional Control. Cell. 2017;169: 13‒23. doi:10.1016/j.cell.2017.02.007

9. Boija A, Klein IA, Sabari BR, Dall’Agnese A, Coffey EL, Zamudio A V., et al. Transcription Factors Activate Genes through the Phase-Separation Capacity of Their Activation Domains. Cell. 2018;175: 1842–1855.e16. doi:10.1016/j.cell.2018.10.042

10. Takada S. Coarse-grained molecular simulations of large biomolecules. Curr Opin Struct Biol. 2012;22: 130‒137. doi:10.1016/j.sbi.2012.01.010

11. Dignon GL, Zheng W, Kim YC, Best RB, Mittal J. Sequence determinants of protein phase behavior from a coarse-grained model. Ofran Y, editor. PLOS Comput Biol. 2018;14: e1005941. doi:10.1371/journal.pcbi.1005941

12. Dannenhoffer-Lafage T, Best RB. A Data-Driven Hydrophobicity Scale for Predicting Liquid‒Liquid Phase Separation of Proteins. J Phys Chem B. 2021;125: 4046‒4056. doi:10.1021/acs.jpcb.0c11479

13. Benayad Z, von Bülow S, Stelzl LS, Hummer G. Simulation of FUS Protein Condensates with an Adapted Coarse-Grained Model. J Chem Theory Comput. 2021;17: 525‒537. doi:10.1021/acs.jctc.0c01064

14. Joseph JA, Reinhardt A, Aguirre A, Chew PY, Russell KO, Espinosa JR, et al. Physics-driven coarse-grained model for biomolecular phase separation with near-quantitative accuracy. Nat Comput Sci. 2021;1: 732‒743. doi:10.1038/s43588-021-00155-3

15. Takada S. Gō model revisited. Biophys Physicobiology. 2015;12: 13‒20. doi:10.2142/biophysico.16.0

16. Kenzaki H, Koga N, Hori N, Kanada R, Li W, Okazaki KI, et al. CafeMol: A coarse-grained biomolecular simulator for simulating proteins at work. J Chem Theory Comput. 2011;7: 1979‒1989. doi:10.1021/ct2001045

17. Tesei G, Schulze TK, Crehuet R, Lindorff-Larsen K. Accurate model of liquid-liquid phase behavior of intrinsically disordered proteins from optimization of single-chain properties. Proc Natl Acad Sci U S A. 2021;118. doi:10.1073/pnas.2111696118

18. Regy RM, Thompson J, Kim YC, Mittal J. Improved coarse-grained model for studying sequence dependent phase separation of disordered proteins. Protein Sci. 2021;30: 1371‒1379. doi:10.1002/pro.4094

19. Niwa H, Ogawa K, Shimosato D, Adachi K. A parallel circuit of LIF signalling pathways maintains pluripotency of mouse ES cells. Nature. 2009;460: 118‒122. doi:10.1038/nature08113

20. Huang D, Wang L, Duan J, Huang C, Tian X, Zhang M, et al. LIF-Activated Jak signaling determines Esrrb expression during late-stage reprogramming. Biol Open. 2018;7: 2‒8. doi:10.1242/bio.029264

21. Chambers I, Colby D, Robertson M, Nichols J, Lee S, Tweedie S, et al. Functional expression cloning of Nanog, a pluripotency sustaining factor in embryonic stem cells. Cell. 2003;113: 643‒655. doi:10.1016/S0092-8674(03)00392-1

22. Mitsui K, Tokuzawa Y, Itoh H, Segawa K, Murakami M, Takahashi K, et al. The homeoprotein nanog is required for maintenance of pluripotency in mouse epiblast and ES cells. Cell. 2003;113: 631‒642. doi:10.1016/S0092-8674(03)00393-3

23. Najafzadeh B, Asadzadeh Z, Motafakker Azad R, Mokhtarzadeh A, Baghbanzadeh A, Alemohammad H, et al. The oncogenic potential of NANOG: An important cancer induction mediator. J Cell Physiol. 2021;236: 2443‒2458. doi:10.1002/jcp.30063

24. Pan GJ, Pei DQ. Identification of two distinct transactivation domains in the pluripotency sustaining factor nanog. Cell Res. 2003;13: 499‒502. doi:10.1038/sj.cr.7290193

25. Mullin NP, Yates A, Rowe AJ, Nijmeijer B, Colby D, Barlow PN, et al. The pluripotency rheostat Nanog functions as a dimer. Biochem J. 2008;411: 227‒231. doi:10.1042/BJ20080134

26. Wang J, Levasseur DN, Orkin SH. Requirement of Nanog dimerization for stem cell self-renewal and pluripotency. Proc Natl Acad Sci. 2008;105: 6326‒6331. doi:10.1073/pnas.0802288105

27. Mullin NP, Gagliardi A, Khoa LTP, Colby D, Hall-Ponsele E, Rowe AJ, et al. Distinct Contributions of Tryptophan Residues within the Dimerization Domain to Nanog Function. J Mol Biol. 2017;429: 1544‒1553. doi:10.1016/j.jmb.2016.12.001

28. Choi K-J, Quan MD, Qi C, Lee J-H, Tsoi PS, Zahabiyon M, et al. NANOG prion-like assembly mediates DNA bridging to facilitate chromatin reorganization and activation of pluripotency. Nat Cell Biol. 2022;24: 737‒747. doi:10.1038/s41556-022-00896-x

29. Krainer G, Welsh TJ, Joseph JA, Espinosa JR, Wittmann S, de Csilléry E, et al. Reentrant liquid condensate phase of proteins is stabilized by hydrophobic and non-ionic interactions. Nat Commun. 2021;12: 1‒14. doi:10.1038/s41467-021-21181-9

30. Chevalier Y, Zemb T. The structure of micelles and microemulsions. Reports Prog Phys. 1990;53: 279‒371. doi:10.1088/0034-4885/53/3/002

31. Saunders A, Li D, Faiola F, Huang X, Fidalgo M, Guallar D, et al. Context-Dependent Functions of NANOG Phosphorylation in Pluripotency and Reprogramming. Stem Cell Reports. 2017;8: 1115‒1123. doi:10.1016/j.stemcr.2017.03.023

32. Mullin NP, Varghese J, Colby D, Richardson JM, Findlay GM, Chambers I. Phosphorylation of NANOG by casein kinase I regulates embryonic stem cell self-renewal. FEBS Lett. 2021;595: 14‒25. doi:10.1002/1873-3468.13969

33. Monahan Z, Ryan VH, Janke AM, Burke KA, Rhoads SN, Zerze GH, et al. Phosphorylation of the FUS low-complexity domain disrupts phase separation, aggregation, and toxicity. EMBO J. 2017;36: 2951‒2967. doi:10.15252/embj.201696394

34. Brumbaugh J, Russell JD, Yu P, Westphall MS, Coon JJ, Thomson JA. NANOG Is Multiply Phosphorylated and Directly Modified by ERK2 and CDK1 In Vitro. Stem Cell Reports. 2014;2: 18‒25. doi:10.1016/j.stemcr.2013.12.005

35. Wang J, Yu H, Ma Q, Zeng P, Wu D, Hou Y, et al. Phase separation of OCT4 controls TAD reorganization to promote cell fate transitions. Cell Stem Cell. 2021; 1‒16. doi:10.1016/j.stem.2021.04.023

36. Henninger JE, Oksuz O, Shrinivas K, Sagi I, LeRoy G, Zheng MM, et al. RNA-Mediated Feedback Control of Transcriptional Condensates. Cell. 2021;184: 207–225.e24. doi:10.1016/j.cell.2020.11.030

37. Chen J, Zhang Z, Li L, Chen BC, Revyakin A, Hajj B, et al. Single-molecule dynamics of enhanceosome assembly in embryonic stem cells. Cell. 2014;156: 1274‒1285. doi:10.1016/j.cell.2014.01.062

38. Xie L, Torigoe SE, Xiao J, Mai DH, Li L, Davis FP, et al. A dynamic interplay of enhancer elements regulates Klf4 expression in naïve pluripotency. Genes Dev. 2017;31: 1795‒1808. doi:10.1101/gad.303321.117

39. Verneri P, Vazquez Echegaray C, Oses C, Stortz M, Guberman A, Levi V. Dynamical reorganization of the pluripotency transcription factors Oct4 and Sox2 during early differentiation of embryonic stem cells. Sci Rep. 2020;10: 1‒12. doi:10.1038/s41598-020-62235-0

40. Takada S, Kanada R, Tan C, Terakawa T, Li W, Kenzaki H. Modeling Structural Dynamics of Biomolecular Complexes by Coarse-Grained Molecular Simulations. Acc Chem Res. 2015;48: 3026–3035. doi:10.1021/acs.accounts.5b00338

41. Li W, Wang W, Takada S. Energy landscape views for interplays among folding, binding, and allostery of calmodulin domains. Proc Natl Acad Sci U S A. 2014;111: 10550‒10555. doi:10.1073/pnas.1402768111

42. Hayashi Y, Caboni L, Das D, Yumoto F, Clayton T, Deller MC, et al. Structure-based discovery of NANOG variant with enhanced properties to promote self-renewal and reprogramming of pluripotent stem cells. Proc Natl Acad Sci U S A. 2015;112: 4666‒4671. doi:10.1073/pnas.1502855112

43. Murthy AC, Dignon GL, Kan Y, Zerze GH, Parekh SH, Mittal J, et al. Molecular interactions underlying liquid−liquid phase separation of the FUS low-complexity domain. Nat Struct Mol Biol. 2019;26: 637‒648. doi:10.1038/s41594-019-0250-x

44. Schuster BS, Dignon GL, Tang WS, Kelley FM, Ranganath AK, Jahnke CN, et al. Identifying sequence perturbations to an intrinsically disordered protein that determine its phase-separation behavior. Proc Natl Acad Sci U S A. 2020;117. doi:10.1073/pnas.2000223117

45. Conicella AE, Dignon GL, Zerze GH, Schmidt HB, D’Ordine AM, Kim YC, et al. TDP-43 α-helical structure tunes liquid‒liquid phase separation and function. Proc Natl Acad Sci U S A. 2020;117: 5883‒ 5894. doi:10.1073/pnas.1912055117

46. Freeman GS, Hinckley DM, Lequieu JP, Whitmer JK, De Pablo JJ. Coarse-grained modeling of DNA curvature. J Chem Phys. 2014;141. doi:10.1063/1.4897649

47. Tan C, Terakawa T, Takada S. Dynamic Coupling among Protein Binding, Sliding, and DNA Bending Revealed by Molecular Dynamics. J Am Chem Soc. 2016;138: 8512‒8522. doi:10.1021/jacs.6b03729

48. Niina T, Brandani GB, Tan C, Takada S. Sequence-dependent nucleosome sliding in rotation-coupled and uncoupled modes revealed by molecular simulations. PLoS Comput Biol. 2017;13: 1‒22. doi:10.1371/journal.pcbi.1005880

49. Tan C, Takada S. Nucleosome allostery in pioneer transcription factor binding. Proc Natl Acad Sci U S A. 2020;117: 20586‒20596. doi:10.1073/pnas.2005500117

50. Jung J, Mori T, Kobayashi C, Matsunaga Y, Yoda T, Feig M, et al. GENESIS: A hybrid-parallel and multi-scale molecular dynamics simulator with enhanced sampling algorithms for biomolecular and cellular simulations. Wiley Interdiscip Rev Comput Mol Sci. 2015;5: 310‒323. doi:10.1002/wcms.1220

51. Kobayashi C, Jung J, Matsunaga Y, Mori T, Ando T, Tamura K, et al. Genesis 1.1: A hybrid-parallel molecular dynamics simulator with enhanced sampling algorithms on multiple computational platforms. J Comput Chem. 2017;38: 2193‒2206. doi:10.1002/jcc.24874

52. Tan C, Jung J, Kobayashi C, Torre DU La, Takada S, Sugita Y. Implementation of residue-level coarsegrained models in GENESIS for large-scale molecular dynamics simulations. PLoS Comput Biol. 2022;18: 1‒30. doi:10.1371/journal.pcbi.1009578

53. Michaud-Agrawal N, Denning EJ, Woolf TB, Beckstein O. MDAnalysis: A toolkit for the analysis of molecular dynamics simulations. J Comput Chem. 2011;32: 2319‒2327. doi:10.1002/jcc.21787

54. Gowers R, Linke M, Barnoud J, Reddy T, Melo M, Seyler S, et al. MDAnalysis: A Python Package for the Rapid Analysis of Molecular Dynamics Simulations. Proceedings of the 15th Python in Science Conference. 2016. pp. 98‒105. doi:10.25080/Majora-629e541a-00e

55. MacKenzie GA, Pawley GS. A neutron scattering study of poly(ethylene glycol) in electrolyte solutions. Macromolecules. 1995;12: 2715‒2735. doi:10.1088/0022-3719/12/14/011

56. Roovers J, Martin JE. The hard-sphere model for linear and regular star polybutadienes. J Polym Sci Part B Polym Phys. 1989;27: 2513‒2524. doi:10.1002/polb.1989.090271209

57. Popielec A, Ostrowska N, Wojciechowska M, Feig M, Trylska J. Crowded environment affects the activity and inhibition of the NS3/4A protease. Biochimie. 2020;176: 169‒180. doi:10.1016/j.biochi.2020.07.009

